# Automatic determination of the handedness of Single-Particle maps of macromolecules solved by CryoEM

**DOI:** 10.1101/2022.03.01.482513

**Authors:** J. Garcia Condado, A. Muñoz-Barrutia, C.O.S. Sorzano

**Affiliations:** Biocruces Bizkaia Instituto Investigación Sanitaria, Cruces Plaza, 48903 Barakaldo, Bizkaia, Spain; Universidad Carlos III de Madrid, Avda. de la Universidad 30, 28911, Leganés, Madrid, Spain; Centro Nacional de Biotecnologia (CNB-CSIC), Darwin, 3, Campus Universidad Autonoma, 28049 Cantoblanco, Madrid, Spain

**Author notes:** Corresponding authors, *Email address:* (C.O.S. Sorzano ^*c*^).

**Keywords:** Electron microscopy, Single Particle Analysis, Validation

## Abstract

Single-Particle Analysis by Cryo-Electron Microscopy is a well-established technique to elucidate the three-dimensional (3D) structure of biological macromolecules. The orientation of the acquired projection images must be initially estimated without any reference to the final structure. In this step, algorithms may find a mirrored version of all the orientations resulting in a mirrored 3D map. It is as compatible with the acquired images as its unmirrored version from the image processing point of view, only that it is not biologically plausible.

In this article, we introduce HaPi (Handedness Pipeline), the first method to automatically determine the hand of electron density maps of macromolecules solved by CryoEM. HaPi is built by training two 3D convolutional neural networks. The first determines α-helices in a map, and the second determines whether the α-helix is left-handed or right-handed. A consensus strategy defines the overall map hand. The pipeline is trained on simulated and experimental data. The handedness can be detected only for maps whose resolution is better than 5Å. HaPi can identify the hand in 89% of new simulated maps correctly. Moreover, we evaluated all the maps deposited at the Electron Microscopy Data Bank and 11 structures uploaded with the incorrect hand were identified.

## 1. Introduction

Single-Particle Analysis is an ill-posed problem because the reconstruction of 3D macromolecular structures from 2D images is not well determined. If all particle image orientations are mirrored, a map is reconstructed that is equally consistent with the measured data but non-superimposable over the map previously reconstructed, see Figure 1. As proteins have a specific handedness [3] only one of the two possible reconstructed maps is the correct reconstruction of the structure.

**Figure 1:**
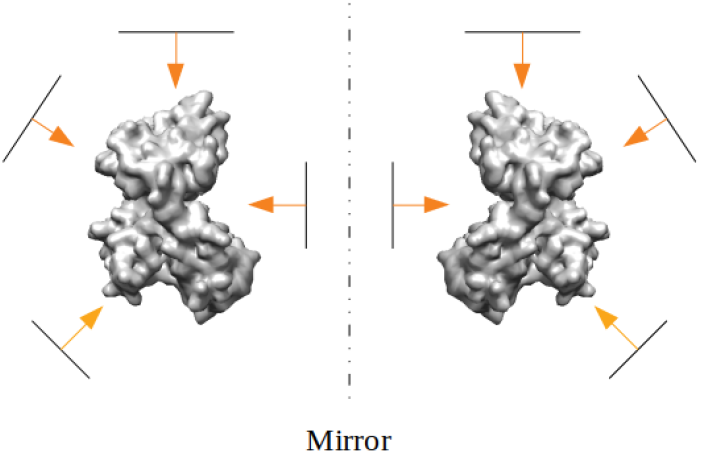
Reconstruction of a structure from the same set of images but with mirror orientations assigned to each image which produces a mirrored version of the structure.

Currently, a trained biologist is required to look at the α-helices rotation to assess the handedness of the map. If incorrect, the reconstructed map is mirrored. The direction of rotation is easily determined at very high resolutions of 1Å but can be difficult at lower resolutions even for experts, see Figure 2. As the resolution decreases, the α-helix slowly transitions from a helix to a cylinder, which no longer has a hand (see Figure 3). Hence, we propose HaPi (Handedness Pipeline) to automatically determine the hand of reconstructed maps using deep learning for resolutions of up to 5Å.

**Figure 2:**
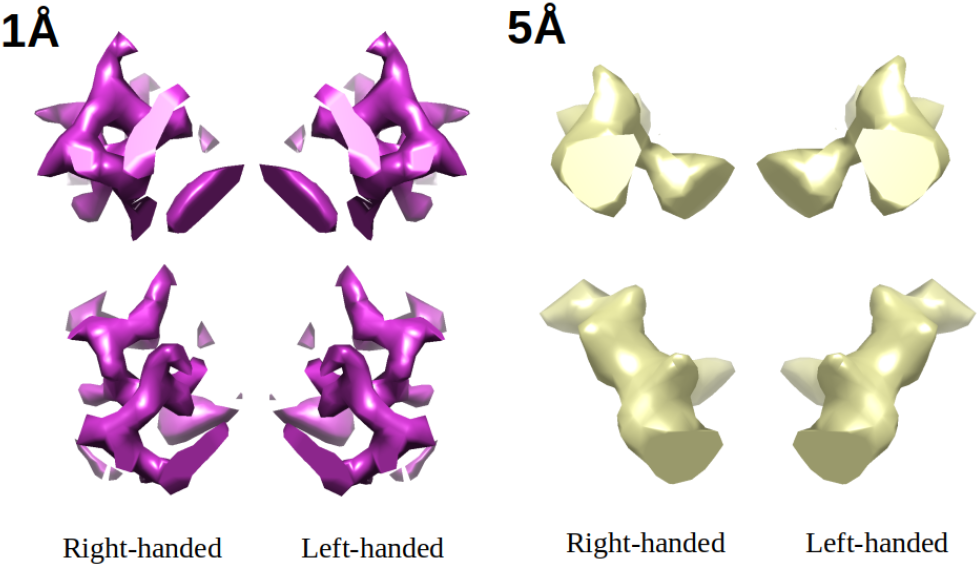
A portion of the same α-helix at 1Å and 5Å with its true structure (right-handed) and mirrored version (left-handed) from different viewing angles.

**Figure 3:**
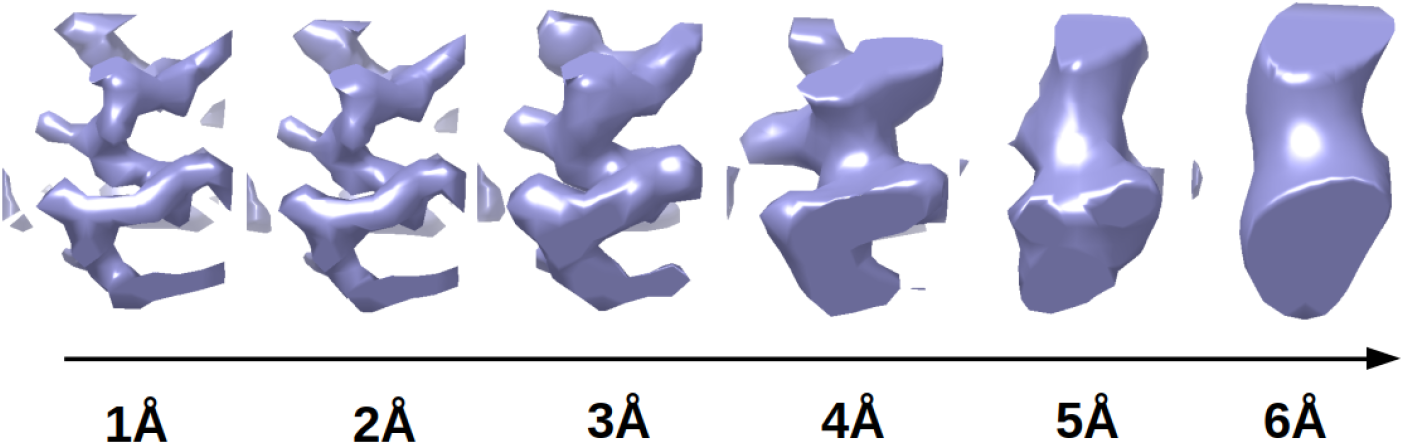
Same α-helix with same viewing angle at different resolutions that shows transition from helical to cylindrical structure.

To the best of our knowledge, there are no algorithms to detect the hand of reconstructed CryoEM maps automatically. The proposed model identifies Secondary Structure Elements (SSE) of interest in the volume and then uses these to detect the hand. There are several previous approaches to automatically determine SSE in electron density maps based on non-machine learning methods [1, 2, 10, 16], machine learning methods [8, 11] and more recently, deep learning techniques [5, 6, 15, 4]. As the latter have shown better performance, in this work, 3D Convolutional Neural Networks (CNNs) are used to determine SSE of interest and detect the hand of a map from small boxes extracted from the map at the location of the SSE.

The number of reconstructed structures is quickly increasing as CryoEM is being widely adopted. HaPi is a valuable tool to guarantee the correctness of automatic image processing pipelines and as quality control in public databases like Electron Microscopy Data Bank (EMDB).

## 2. Methods

The HaPi package is freely available to use and documented via GitHub (https://github.com/JGarciaCondado/EMapHandedness). All the code for the methods described can be found in the same link. HaPi is also available in Scipion [7] and Xmipp [12].

### 2.1. Pipeline

HaPi determines the hand of electron density maps, Figure 4 considering as inputs a Coulomb Potential map (*V_f_*) and a mask of the non-background voxels (*V_mask_*). Maps are first preprocessed by resampling to 1Å/voxel and low-pass filtering to 5Å to match the training data. This procedure reduces noise and homogenizes local resolutions.

**Figure 4:**
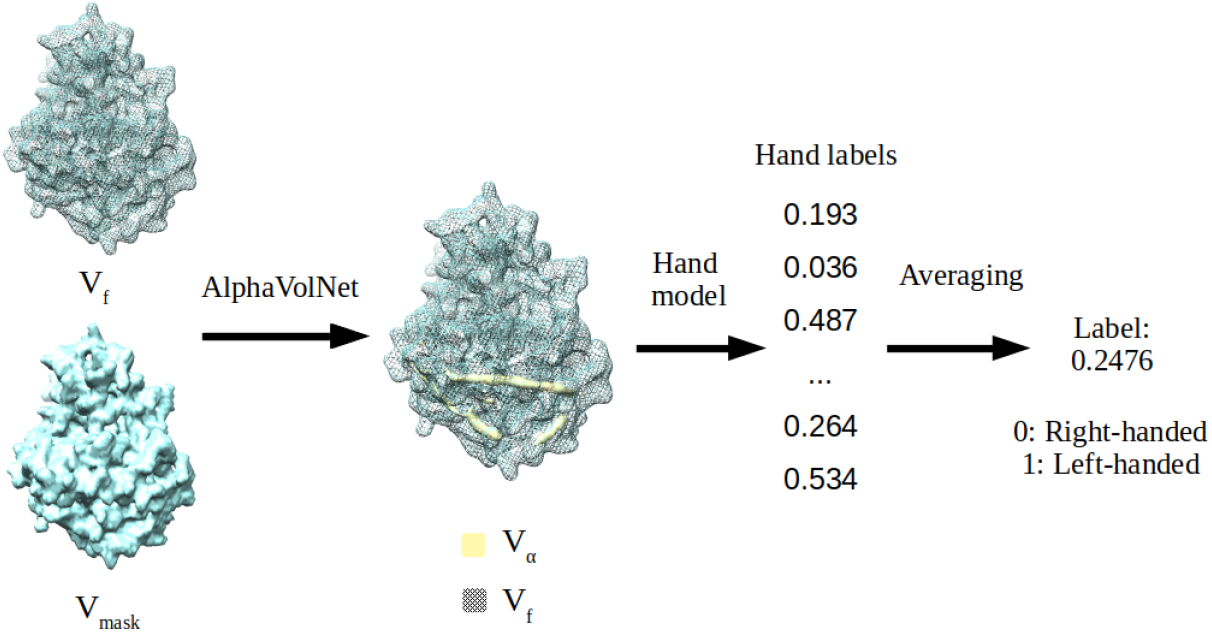
Diagram of the HaPi pipeline with the inputs and outputs at the different stages and the models used to generate each. The input to HaPi is *V_f_* (a Coulomb potential map) and *V_mask_* (a mask of the overall structure). These are first passed through AlphaVolNet, which outputs a mask *V_α_* of the proposed location of α-helices (yellow). Boxes are extracted from *V_f_* at the location of voxels in *V_α_* that contain an α-helix. They are passed through the hand model to be given a hand value. Values are averaged to assign a value to the overall map. The pipeline has 208,163 parameters.

AlphaVolNet determines the location of α-helices in the whole volume. It takes as input *V_f_* and *V_mask_* and outputs *V_α_*, which is a mask containing the location of the α-helices found. It does so by taking at each non-background voxel location of *V_f_* a box of dimensions 11 × 11 × 11 voxels and passed through the trained 3D CNN α-SSE model. Then, if the label is above a threshold *t_α_, V_α_* is set to true at that location.

HandNet is used to determine the hand of a map from *V_α_* and *V_f_*. At each active voxel of *V_α_*, a box of dimension 11 × 11 × 11 voxels is extracted at that location from *V_f_* and passed through the trained hand model. Then, a hand value is given to the map by consensus of all the labels of the boxes. The consensus value is the average of all hand predictions of each α-box.

### 2.2. 3D CNN model

The same 3D CNN model is used for the SSE and hand determination task. A 3D CNN is an extension of 2D CNNs that deals with volumes instead of images. The whole architecture of the 3D CNN design can be seen in Figure 5. All 3D convolutional layers and the first connected layer are followed by ReLu activation functions. A sigmoid function follows the last fully connected layer. The 3D CNN has in total 104,080 parameters.

**Figure 5:**
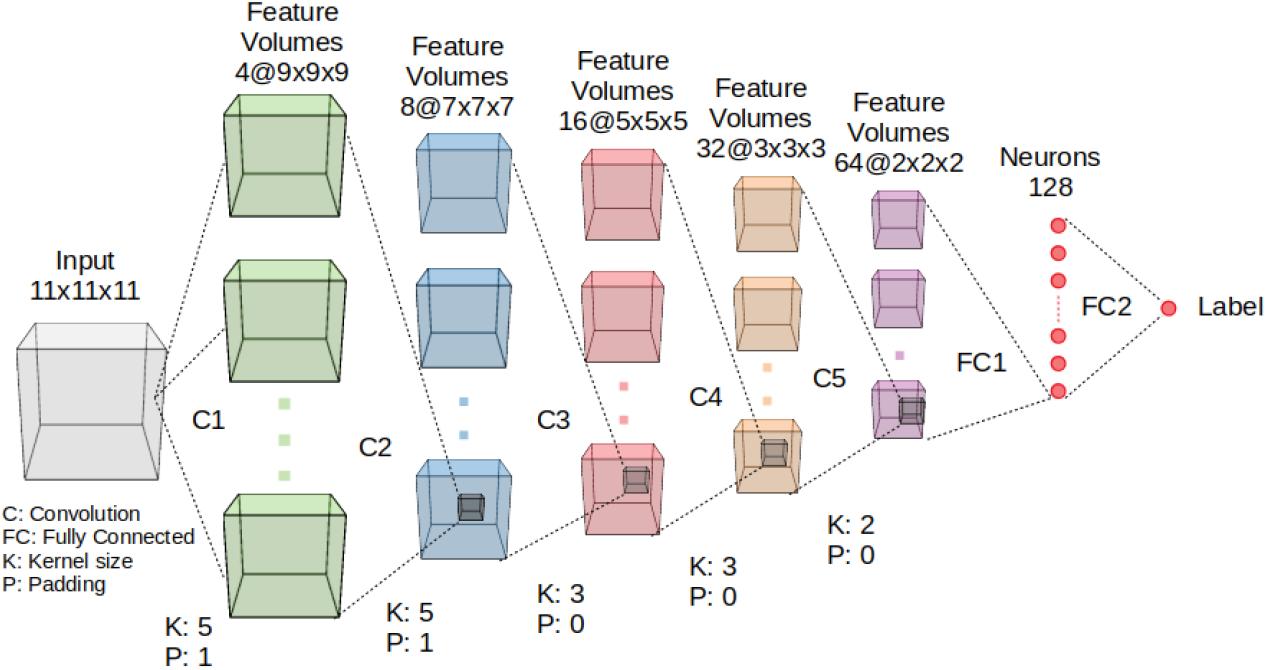
3D CNN model diagram used for both Secondary Structure Element (SSE) determination and hand determination. The input to the network is a box containing the electron density at each voxel normalised to the range 0 to 1. The output of the network is a value between 0 and 1. In the SSE determination task, a label of 1 represents that the SSE of interest is found within the box, and 0 means it is not. In the hand determination task, a label of 1 means the box is left-handed, and 0 means it is right-handed.

Input boxes are preprocessed by clipping all negative values to 0 and rescaling the box to be in the range 0 and 1.

The training data differs for each task. For the SSE determination task, 261 experimental maps and their respective fitted atomic models were downloaded from EMDB [14]. In total, 9,985 boxes of dimension 11 × 11 × 11 voxels were extracted from the centroids of alpha helices in the structures and another 9,985 boxes of the same size from random parts of the structure. For the hand determination task, the Xmipp library [12] was used to simulate Coulomb potential maps of 12,343 atomic models using Electron Atomic Scattering Factors [13]. From these, 101,944 boxes of dimension 11 × 11 × 11 voxels from the center of α-helices were extracted to use for training. Half of the boxes were randomly flipped to obtain left-hand helices.

The training strategy is similar for both tasks. For the task of SSE determination, a label of 1 is given if it contains the SSE of interest and 0 if it does not. For the hand determination task, a label of 1 is given if the SSE is left-handed (this is the hand that is seldom found in nature) and a label of 0 if it is right-handed. A binary cross-entropy loss function is used for training with an Adam optimiser, a learning rate of 0.001 and batches of size 2,048. The model is trained for 50 epochs. An early stopping strategy is adopted where the model saved at the epoch where the validation set loss stagnates is used as the final model.

Weight initialisation differs for the SSE and hand models. For SSE determination, the model has been initialised with the default PyTorch settings for weight initialisation. The hand model was first trained on 1Å data and initialised with the default PyTorch settings. Then, a transfer learning approach is used to train on 5Å data for hand determination by using the weights of the model trained on 1Å on initialization.

## 3. Results

The training and validation curves for each of the models trained can be seen in Figure 6. The network is unable to learn how to identify the hand of boxes at 5Å resolution without transfer learning as seen in the high loss value in Figure 6. When the weights are initialized at 1Å the loss value significantly decreases during training. The validation set loss stagnates for the models trained after only 15 epochs.

**Figure 6:**
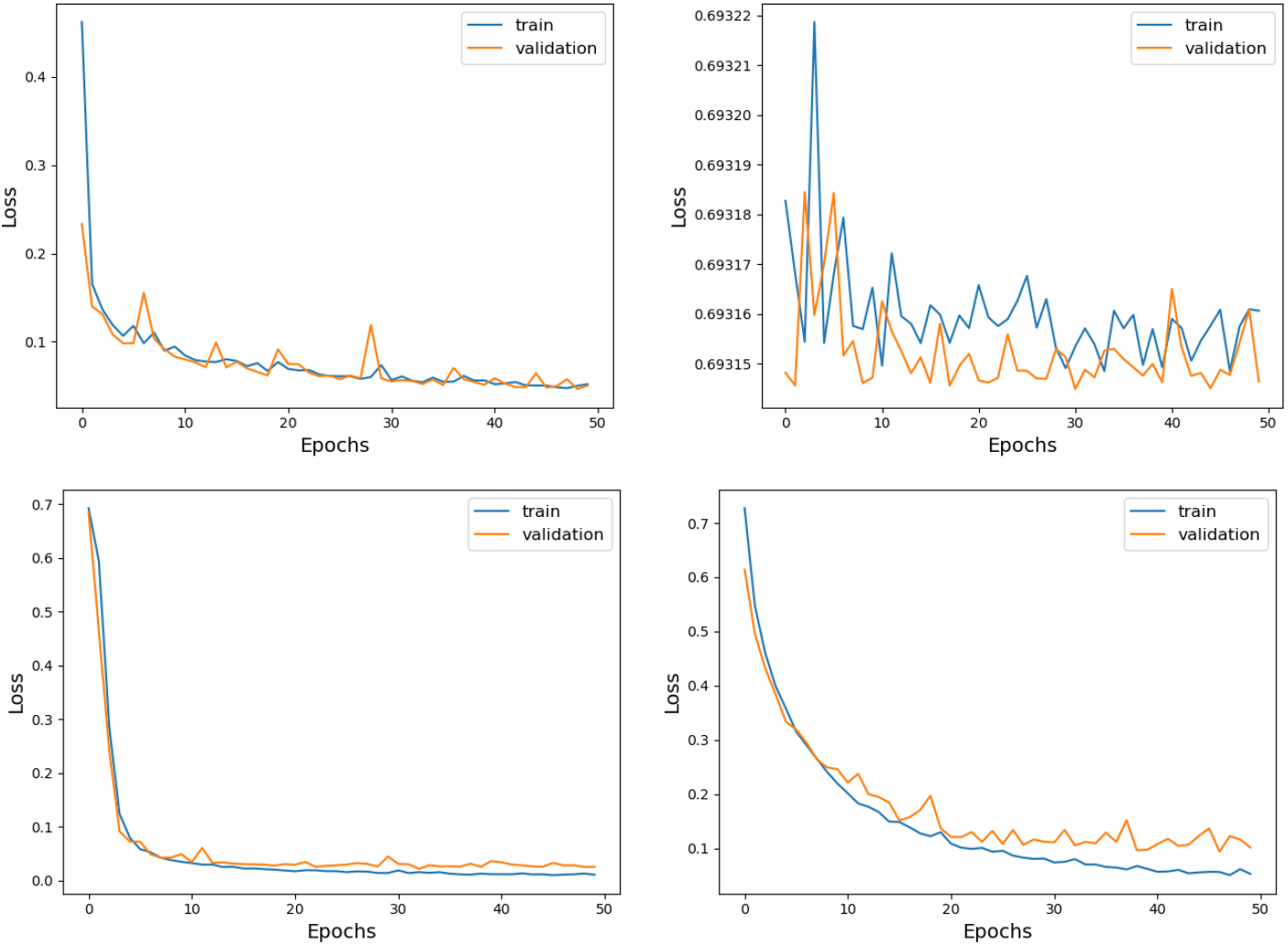
Training and validation loss for each of the 3D CNN models trained. (Top left) Training and validation loss for α-helix determination at 5Å. (Top right) Training and validation loss for hand determination at 5Å. Clearly the network is not able to generalize as the loss stagnates at a high value and is unable to correctly determine the hand. (Bottom left) Training and validation loss for hand determination at 1JI. (Bottom right) Training and validation loss for hand determination at 5Å with weights initialized at those of the model trained at 1Å. With transfer learning strategies the model is able to learn how to correctly classify the hand at lower resolution.

A dataset consisting of 3,119 atomic models was used to simulate maps at 5Å. These were split into validation and test sets of 30% and 70%, respectively. The validation set was used to set *t_α_*, which controls the stringency of the model to accept voxels as containing an α-helix. It was set to *t_α_* = 0.7 to maximize hand accuracy in the validation set. HaPi is able to identify the hand in 89% of new simulated maps correctly. The resulting hand prediction values for the test set is shown in Figure 7.

**Figure 7:**
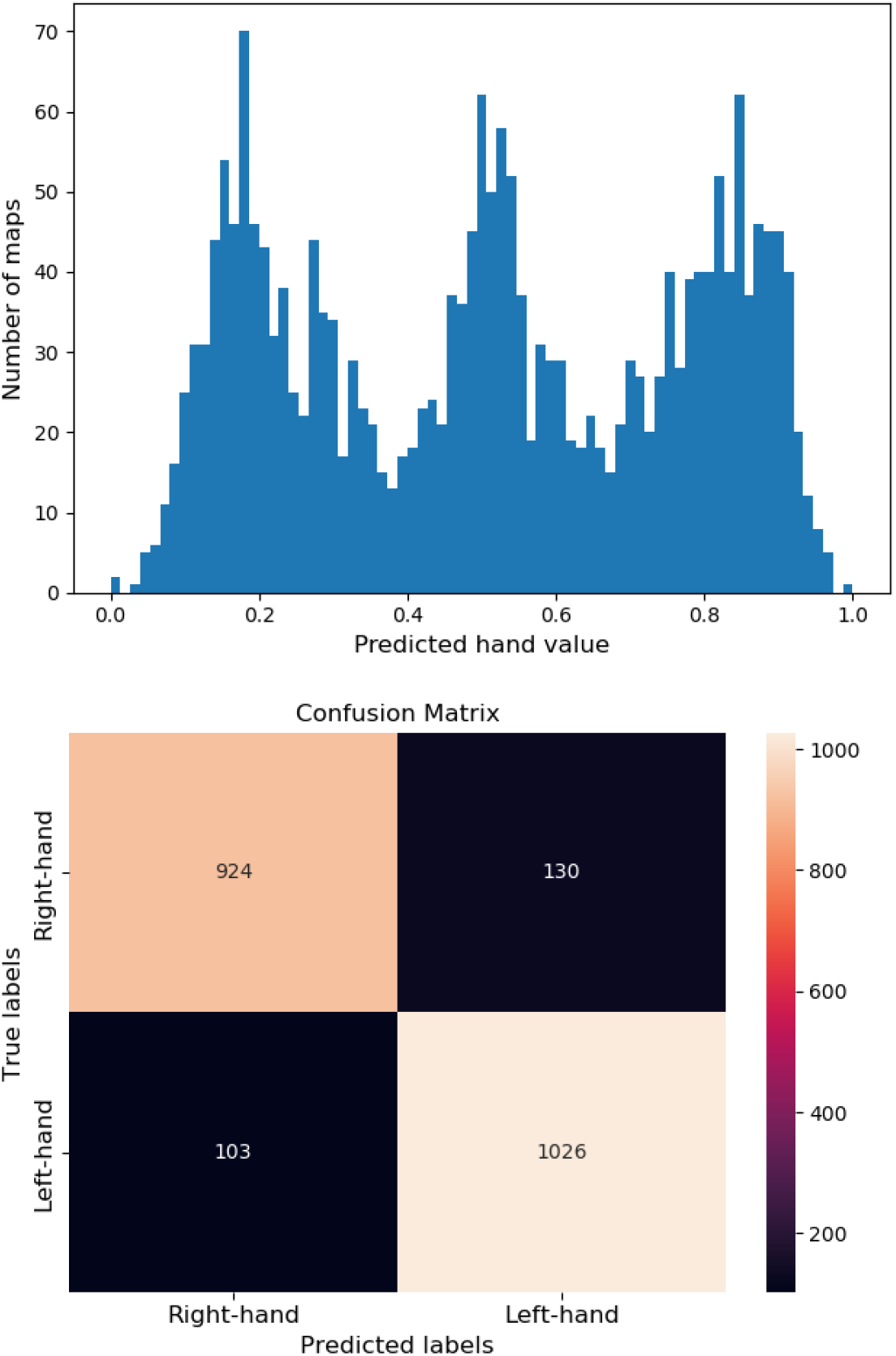
Individual predicted hand values for each simulated map (5Å resolution) in test set (above) and confusion matrix (below) for *t_α_* = 0.7. *t_α_* controls the stringency of the model to accept voxels as containing an α-helix. Simulated maps were left as is to be right-handed or mirrored to be left-handed. Each map was then passed through the HaPi pipeline to receive a hand value. Maps with values above 0.5 were given a left-hand label and maps with values below 0.5 were given a right-hand label

All deposited experimental maps in EMDB with resolution below or at 5Å were downloaded and passed through HaPi. In total, 8,061 maps were downloaded. Maps deposited in EMDB should all be right-handed. The resulting hand predictions for structures can be seen in Figure 8. All structures with a high chance of being left-handed (those with hand value > 0.6, which are 285 structures) were manually checked to assess if they were left-handed. Eleven of these structures were indeed present in the database despite incorrect handedness (EMD-9890, EMD-10012, EMD-22052, EMD-22053, EMD-22056, EMD-22057, EMD-22058, EMD-11082, EMD-11083, EMD-20213 and EMD-23584). Assuming that all structures with hand value < 0.6 were uploaded to EMDB with correct handedness (right-hand) and the error rate is only 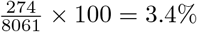.

**Figure 8:**
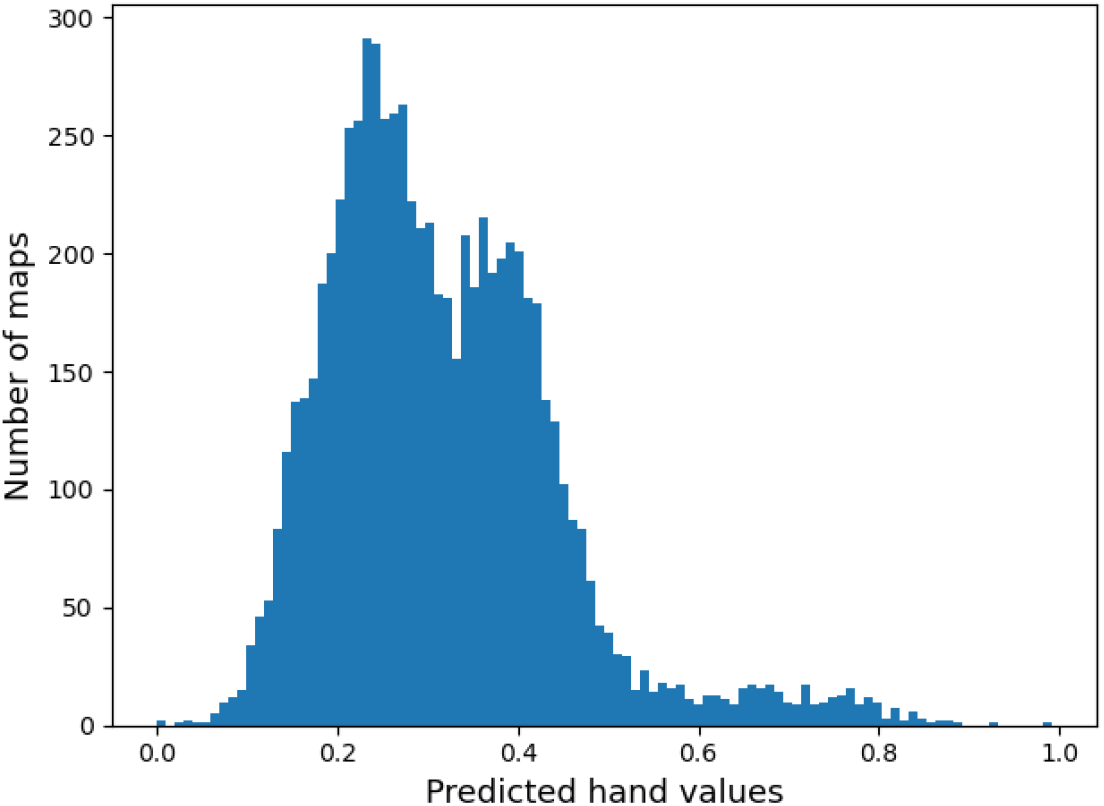
Individual predicted hand values by HaPi for each map in the EMDB database with 5Å or less for *t_α_* = 0.7. *t_α_* controls the stringency of the model to accept voxels as containing an α-helix. Values close to 0 indicate the map is right-handed and values close to 1 indicate the structure is left-handed.

## 4. Discussion

HaPi is able to automatically determine the hand of reconstructed macromolecular structures for intermediate resolutions of 5Å or below. Therefore, HaPi is a valuable tool for validation in databases to avoid incorrect structures to be uploaded. It will also reduce the time biologists dedicate to checking the handedness during image processing and facilitate the construction of automatic image processing pipelines. The method implemented in Scipion automatically return a flipped volume if the hand value is above a threshold set by the user (a value of 0.6 is recommended.) As the number of maps with a resolution of 5Å or below is increasing at a fast pace, this tool can become very useful for many.

HaPi is robust as it has a high accuracy of 96.6% on previously unseen experimental data. Although one of the networks was trained on synthetic data, it can still generalize to experimental data. Some of the downloaded maps contained filaments, electron crystallography data and DNA. This type of data was not used for training. Hence, part of the error rate could be due to including these data during testing. Also, structures that do not contain a considerable number of α-helices are not well predicted as α-helices are the basis for determining the hand.

HaPi is versatile because it does not output a discrete hand label but a value between 0 and 1. Closer to 0 means it is right-handed, and 1 means it is left-handed. Hence, HaPi estimates how sure it is about its decision. Changing the threshold of when to accept a structure as right-handed gives control to the user on stringency. The results on simulated data structures with values lower than 0.4 are highly likely to be right-handed, and those above 0.6 are likely to be left handed. HaPi is unsure of the hand for structures whose hand value is between 0.4 and 0.6, 90% of the structures whose hand was wrongly determined lied between these values.

The structures whose hand values lies between 0.4 and 0.6 and which HaPi has difficulty determining belong to two different groups. The first group would be structures that do not have clear α-helices or have only a few of them. As the algorithm searches for this, its predicting power is diminished if it does not find suitable candidates. The second group are structures whose resolution is very close to 5Å. With lower resolution, there is less information about the hand encoded in the structure, and therefore it is more difficult for HaPi to be sure about its decision.

An experiment was also run to determine which type of SSE was better suited for hand determination between α-helices and β-sheets. At 5Å the 3D CNN trained on α-helices had an accuracy of 95.5% when detecting the hand. On 5Å β-sheets it had an accuracy of 61.0% when detecting the hand; α-helices are better for determining the hand at box level than β-sheets. A biologist can easily distinguish the hand of an α-helix at high resolutions. Even at high resolutions, it is difficult to determine the hand of a β-sheet as it requires looking at the side-chains and their orientation. Hence, the α-helix model was used to build the whole pipeline.

The 3D CNN trained on data at 6Å gave chance level results. The inability of the network to identify the hand at 6Å is the result of the hand information not being available at such resolution. The minimum resolution at which the hand of an α-helix can be determined is that of its pitch. α-helices have an average pitch of 5.4A [9]. At resolutions above 5.4Å, the structure is no longer a helix but a cylinder. Blurred helices become cylinders as information between amino acids below and above a turn merge to form a continuous chain resembling a solid structure rather than a spring coil. If a cylinder is reflected, its mirror version is superimposable over the non-mirrored cylinder. Therefore, a cylinder has no hand.

Being able to determine the hand of an experimental map at or below 5Å is still useful. Structures with resolutions below 5Å already represent more than 50% of all depositions at the EMDB. However, at resolutions below 4Å, the hand can be easily identified by manual inspection of the map as seen in Figure 3. Still, structures between 4Å and 5Å of resolution represent 13.1% of all deposited structures, and this is likely to increase as the resolution of CyroEM maps improves. Hence, this will be a valuable tool to easily and automatically determine the hand at this resolution range.

## 5. Conclusion

HaPi has shown that it is possible to automatically determine the hand of a map without the necessity of inspection by a trained expert. This is even the case for intermediate resolutions where the hand is not clear from visual inspection. HaPi offers a valuable tool for validation in databases and increases biologists’ efficiency by reducing the need for hand inspection during the processing.

The automatic determination of the map hand is a very useful task in the construction of automatic image processing pipelines in CryoEM, and reduces the probability of depositing incorrect maps in public databases such as the EMDB.

## Acknowledgements

This work is supported in part by Ministerio de Ciencia, Innovación y Universidades, Agencia Estatal de Investigación, under grant PID2019-109820RB-I00, MCIN/AEI /10.13039/501100011033/, cofinanced by European Regional Development Fund (ERDF), “A way of making Europe.”

The authors acknowledge the economical support from MICIN to the Instruct Image Processing Center (I2PC) as part of the Spanish participation in Instruct-ERIC, the European Strategic Infrastructure Project (ESFRI) in the area of Structural Biology and Grant PID2019-104757RB-I00 funded by MCIN/AEI/ 10.13039/501100011033/ and “ERDF A way of making Europe”, by the “European Union”. Comunidad Autónoma de Madrid through Grant: S2017/BMD-3817. Instituto de Salud Carlos III: IMP/00019 (ISCIII-SGEFI / ERDF). CSIC and JAE Intro Program: JAEINT_20_01330. European Union (EU) and Horizon 2020 through grant: HighResCells (ERC - 2018 - SyG, Proposal: 810057)

## Notes

https://github.com/JGarciaCondado/EMapHandedness

### Competing Interest Statement

The authors have declared no competing interest.

